# Parallel 1024-ch Cyclic Voltammetry on Monolithic CMOS Electrochemical Detector Array

**DOI:** 10.1101/799916

**Authors:** Kevin A. White, Geoffrey Mulberry, Brian N. Kim

## Abstract

Large-scale microelectrode arrays offers enhanced spatiotemporal resolution in electrophysiology studies.. In this paper, we discuss the design and performance of an electrochemical detector array which is capable of 1024-ch parallel cyclic voltammetry (CV) as well as other electrochemical measurements. The electrochemical detector is fabricated using a custom-designed CMOS chip which integrates both the circuity and on-chip microelectrode array, to operate and record from electrochemical measurements. For parallel 1024-ch recordings, 1024 capacitor-based integrating transimpedance amplifiers (TIA) are designed and integrated. The TIA design features the bipolar capabilities for measuring both negative and positive electrochemical currents due to reduction and oxidation of molecules. The resulted dynamic range of this TIA is −700 pA – 1968 pA. CV can be used to examine the quality of electrochemical electrodes by measuring the double-layer capacitance. Double-layer capacitance forms at the electrode-electrolyte interface and is a function of the effective area of the electrode. Thus, a contaminated electrode can have smaller effective area resulting in smaller double-layer capacitance. Using the parallel CV capability of the monolithic CMOS device, the double layer capacitance of all 1024 electrodes are simultaneously measured to examine the status of the electrodes’ surface in real time. The initial measurement of the electrode array showed a mean capacitance of 466 pF. After plasma treatment to remove contamination on the electrode’s surface, the increased capacitance was 1.36nF nearly tripling the effective surface area. We have successfully developed of 1024-ch electrochemical detector array using the monolithic CMOS sensor. The CV functionality was validated by measuring the double-layer capacitance of the on-chip electrode array. This method can accelerate the characterization of a massive electrode array before analytical experiments to provide well-controlled electrochemical electrodes, which is crucial in conducting reliable electrochemical measurements.

## I. INTRODUCTION

Microelectrode arrays (MEA) have been developed to enhance the spatial resolution of electrochemical recordings, such as mapping the level of H_2_O_2_ in relation to neurological diseases in a brain slice [1], electrochemical imaging of redox molecules diffusing across an MEA [2], and imaging of metabolites in biofilms [3]. To enable parallel recordings, an array of readout circuits is required to interface with MEAs to provide simultaneous amplification and signal processing. Emerging devices for large-scale electrochemical detection are exploring complementary metal-oxide-semiconductor (CMOS) technology to interface with an MEA for high spatial and temporal resolution. Programmable CMOS devices were developed with the capability of performing voltammetry as well as potentiometry [4] or amperometry [5]. Using an array of 100 amperometric amplifiers in a CMOS device, the neurotransmitter release from chromaffin cells were recorded [6], [7].

In this field of evolving technology, the quality of the MEAs must be evaluated after fabrication or repeated use by characterizing the electrodes for possible contamination and its effective surface area. This evaluation step is crucial to perform controlled analytical experiment using the well-characterized electrodes. To characterize the electrodes, cyclic voltammetry (CV) can be used to study various responses of the electrode, such as its impedance as well as sensitivity to oxidation and reduction reactions [8]–[10]. The double-layer capacitance, created by the interface created between an electrode and the electrolyte, can be measured to determine the effective surface area of the electrode-electrolyte interface. In electrochemical measurements, including amperometry and voltammetry, the electrode is held at a potential and oppositely charged ions in the electrolyte (Fig. 1a) will attract to the electrode’s surface and create a capacitive interface (Fig. 1b). When the electrode is contaminated, the effective area to interface with the electrolyte is reduced, diminishing the sensitivity of the electrode. As the contamination is removed from the surface of the electrode, CV will reveal an increase in capacitance through an increased area of the electrode-electrolyte interface (Fig. 1c).

**Fig. 1.**
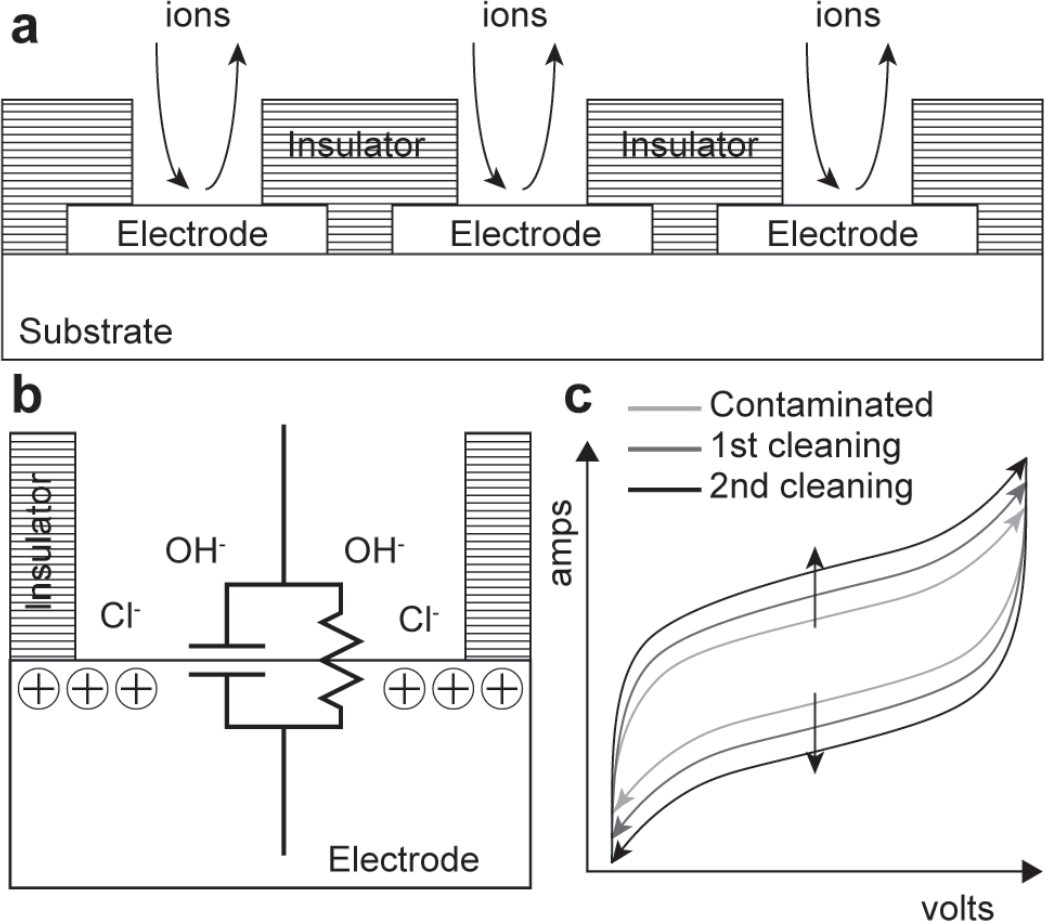
Double-layer capacitance and cyclic voltammetry progression of a contaminated electrode. (a) The electrolytic solution put on the CMOS electrochemical device introduces ions to the electrode. (b) The electrode is held at a potential for electrochemical measurements, attracting oppositely charged ions in the electrolytic solution and creating a double-layer capacitance at the electrode-electrolyte interface. (c) A contaminated electrode exhibits less sensitivity compared to a pristine electrode. As the electrode is cleaned, the progression of the response shows an increase in sensitivity.

Previously, we have presented the development of a monolithic CMOS device with a 1024 on-chip electrode array and 1024 parallel channels for the recording of neurotransmitter release at the single-cell level with a single-vesicle resolution [11]–[13]. Traditional methods of recording single-cell neurotransmitter release at the single-vesicle resolution are low-throughput, costly, and time-consuming. Using the developed device, parallel amperometric recordings of neurotransmitter release were recorded from over 70 human neuroblastoma cells, SH-SY5Y, in a single experiment with over 7000 vesicle release events observed. SH-SY5Y cells are often used as in vitro models to study Parkinson’s disease due to the similarity of their catecholaminergic neuronal properties to dopaminergic neurons in the midbrain [14], [15]. Using the developed high-throughput system, the presented device provides enhanced capabilities to observe secretion from individual vesicles to aid in the study of the molecular level mechanisms of neurological diseases [16], such as Parkinson’s disease, and the sides effects of drug treatments [17], [18].

In this paper, we present the design and performance of the capacitive transimpedance amplifier which has the capability to perform bipolar measurements to enable the characterization of the integrated electrode array using parallel CV. Characterization of the electrode array reveals the state of the surface of individual electrodes, which can often be contaminated and impede effective electrochemical recordings. Also, the effectiveness of plasma treatment towards removing contaminants can be quantified through characterization of the electrode. In this paper, electrode characterization is performed on the monolithic CMOS electrochemical detector array using parallel CV measurements. In Section II, the design and performance of the designed bipolar TIA are discussed as well as the methodology used to integrate the TIA into an electrochemical detector array. In Section III, we discuss the post-CMOS processing used to integrate the electrodes and insulation onto the CMOS chip as well as parallel CV measurements on an electrode array. In Section IV, we discuss the characterization of the integrated electrode array on the monolithic CMOS device using CV before and after a plasma cleaning process.

## II. DESIGN OF BIPOLAR TRANSIMPEDANCE AMPLIFIER

In this section, we detail the design of a bipolar transimpedance amplifier (TIA) and its integration into a 1024 electrochemical detector array.

### A. Bipolar Capacitive Transimpedance Amplifier

To identify oxidation and reduction reactions using CV, bipolar current measurement is required. To take this into account, a TIA with an integrating capacitor is designed to enable bipolar measurement using the DC offset (I_offset_) introduced to the TIA (Fig. 2). With I_offset_ introduced as a baseline measurement, the current from electrochemical reactions at the electrode (I_elect_) can be either positive (+I_elect_) or negative (−I_elect_) due to oxidation or reduction respectively. To obtain a high injection efficiency for the presented TIA, a regulated cascode topology is used [19]. Power consumption and area of the amplifier is minimized by using a simple design using five transistors for the operational amplifier (OPA) [11]. Negative feedback through the cascode transistor regulates the potential of the electrode to be the same as the non-inverting input V_pos_. As the total current (I_int_) is integrated on the integration capacitor (C_int_), the voltage of the readout node V_out_ is reduced. At a periodic interval (Δt), the total voltage drop is readout from the amplifier and C_int_ is reset during readout to begin the next integration cycle. This periodic interval determines the sampling rate of the amplifier. To achieve a 10 kS/s sampling rate, the period readout interval is 100 μs. In the absence of electrochemical reactions at the electrode, only I_offset_ is integrated resulting in a readout of ΔV_offset_ (Fig. 2a). During oxidation reactions, electrons are being released into the electrode, creating a current entering the electrode node which results in an integration current above the offset of I_int_ = I_offset_ + I_elect_. The total integrated current is larger during oxidation, resulting in a larger readout of Δ(V_offset_ + V_elect_) (Fig. 2b). Conversely, in reduction reactions electrons are being taken from the electrode resulting in a total integration current below the offset of I_int_ = I_offset_ – I_elect_. This reduction in integration will result in a smaller readout of Δ(V_offset_ – V_elect_) (Fig. 2c) during reduction reactions. To sustain the negative feedback of the amplifier, I_elect_ can be as low as −I_offset_ before the TIA effectively turns off. The TIA’s transimpedance gain is determined by both C_int_, estimated to be 116 fF, and Δt. Knowing the transimpedance gain, the bipolar dynamic range of the TIA is −I_offset_ to + (C_int_×(V_DD_−V_pos_)/Δt) – I_offset_. With a V_DD_ of 3.3 V, V_pos_ of 1.0 V, I_offset_ of 700 pA, C_int_ of 116 fF, and Δt of 100 μs, the dynamic range is −700 pA – 1968 pA.

**Fig. 2.**
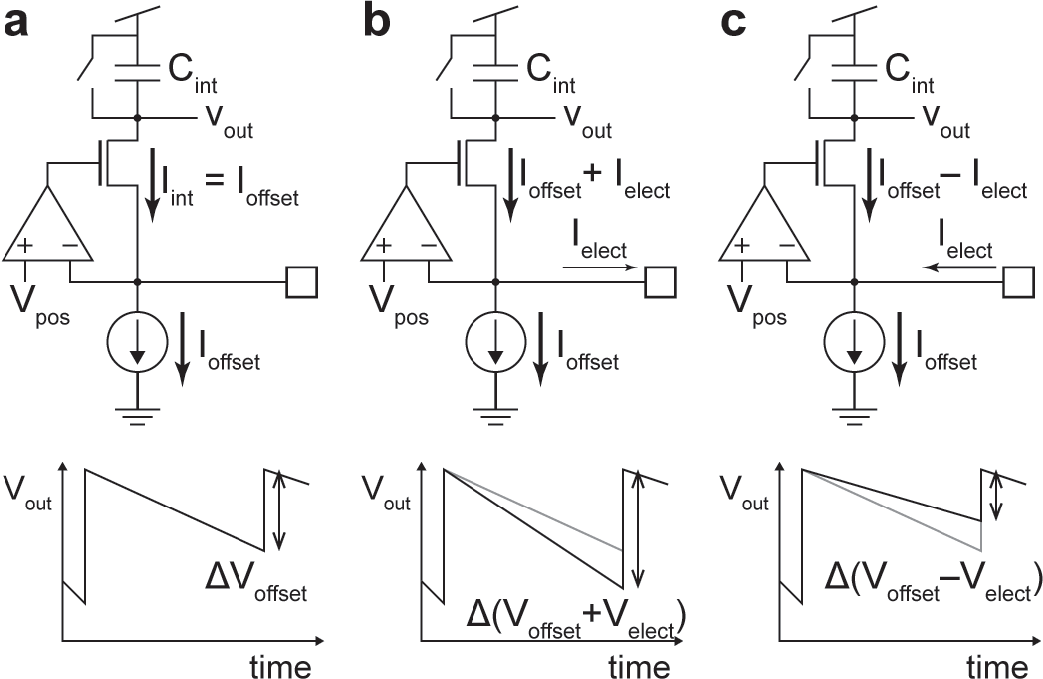
Designed bipolar capacitive TIA. (a) In the absence of electrons transferred to/from the electrode, only I_offset_ integrated and ΔV_offset_ is readout. (b) During oxidation reactions, the current integrated is increased to I_offset_ + I_elect_, increasing the readout to Δ(V_offset_ + V_elect_). (c) During reduction reactions, the current integrated is decreased to I_offset_ − I_elect_, reducing the readout to Δ(V_offset_ − V_elect_).

### B. Bipolar Measurements and Characteristics

To characterize the bipolar functionality of the designed amplifier, a current is injected into the electrode node of the amplifier while I_offset_ remains constant. The current is created by connecting a l-GΩ resistor to the electrode node and applying a DC voltage to the other terminal of the resistor. Because the voltage at the electrode node is regulated through the feedback, the voltage drop across the resistor can be precisely controlled for current injection into the TIA. The linearity of the amplifier’s output is studied by applying various levels of bipolar current. For this experiment the non-inverting input V_pos_, used to set the electrode potential, is 1.0 V. The range of the DC voltage applied is 0.3 V – 1.7 V, generating an input current range of +700 pA – −700 pA. The output of the amplifier, as the voltage applied to the 1-GΩ resistor is changed every 5 seconds, is shown in Fig. 3a and the reference of 0 pA corresponds to the integration of only the I_offset_ current, set to 700 pA. By mapping the output voltage versus the injected current, the linearity of the TIA can be determined as shown in Fig. 3b. The fit shown in the figure gives a transimpedance gain of 0.679 mV/pA with a coefficient of determination, R^2^, of 0.998, indicating a high level of linearity for bipolar current measurement.

**Fig. 3.**
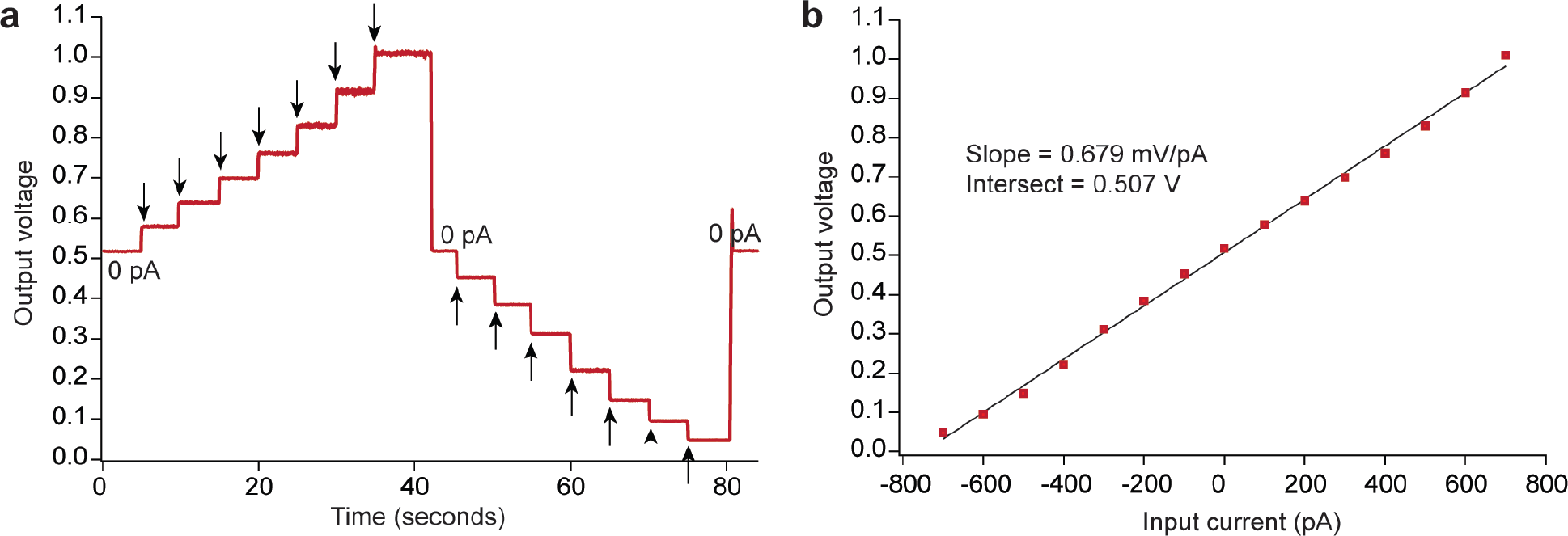
Bipolar current measurement of the designed TIA (a) Voltage readout of the TIA at different levels of current injected into the electrode. (b) Linear curve fitting illustrating the input current vs output voltage linearity. The TIA used exhibits a transimpedance gain of 0.679 mV/pA with a R^2^ value of 0.998, indicating the highly linear performance of the TIA.

Further characterization of the amplifier’s bipolar capabilities is studied by recording alternating signals. Alternating current is injected into the electrode node using a 1-MΩ resistor connected to the electrode node. A 1-MΩ resistor is chosen for this experiment to remove the influence of a zero in the acquired set of measurements. Most component resistors have a parasitic capacitance near 500-fF and thus the zero created by the 1-MΩ resistor is ~318 kHz. The introduction of a zero can increase the amplitude of the input current and distort the results in the low-frequency range. The tested cases are at frequencies of 10, 100, and 1 kHz with each frequency using the same AC amplitude of 200 μV and at super-imposed DC levels of +100pA, 0pA, and −100pA. These sets of measurements are shown in Fig. 4 and they validate the amplifier’s ability to measure bipolar alternating current. The experiments performed confirm the capability of the designed capacitive TIA to measure bipolar current.

**Fig. 4.**
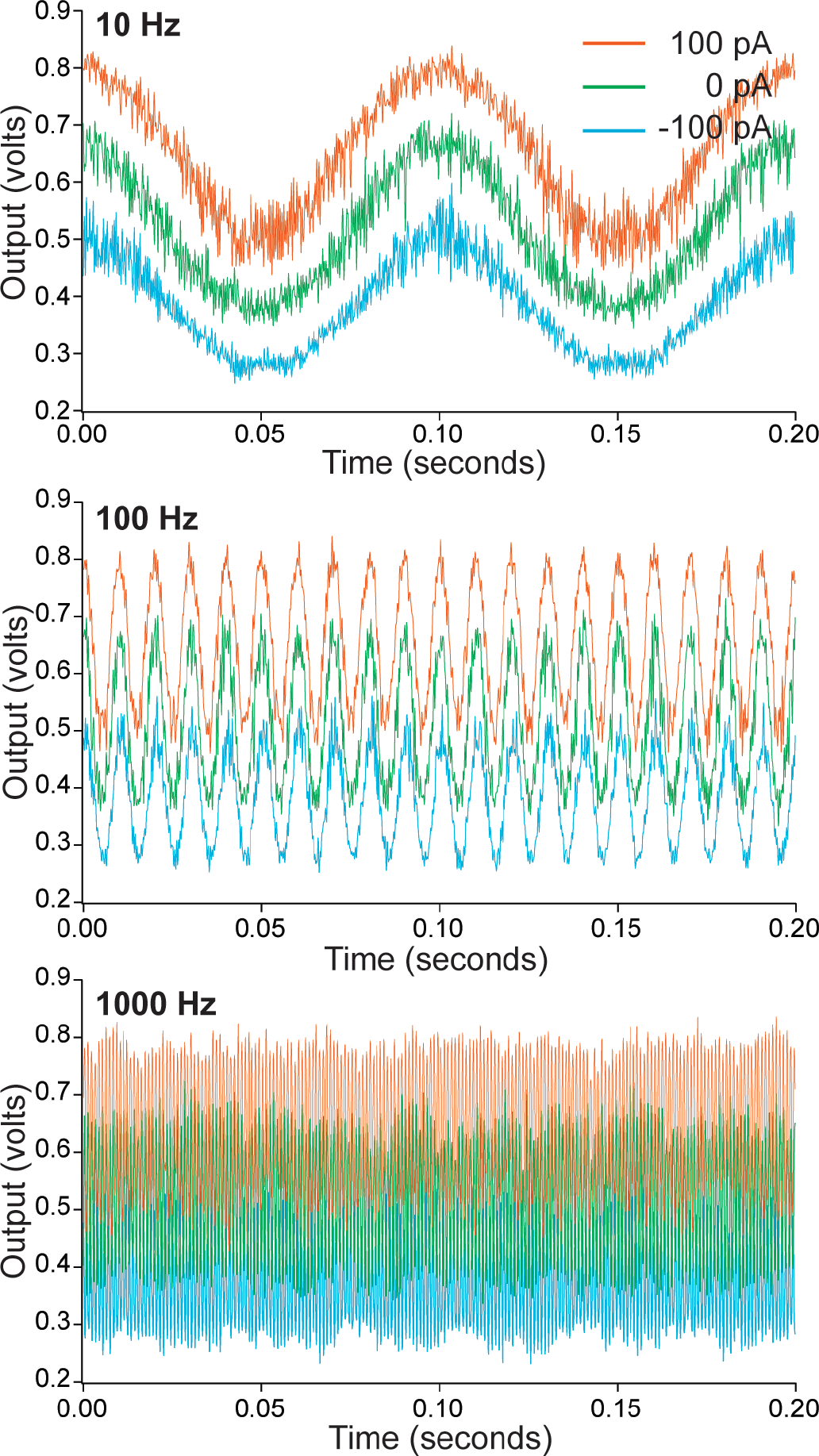
Measurement of alternating current at different levels of DC injection and at frequencies of 10, 100, and 1000 Hz. Under the various DC injection cases, the injected AC signal is evident and similar between the various input frequencies.

### C. Integration of the Electrochemical Detector Array

To integrate an array of amplifiers for parallel electrochemical measurements, a half-shared OPA structure is used, which has been previously studied [11], [12]. The purpose of the half-shared OPA design is to reduce the area required to create a dense sensor array by sharing one non-inverting input with multiple inverting inputs (Fig 5). In the presented device, one non-inverting half of an OPA is shared by groups of 4 TIAs, each of which occupies 30 × 30 μm^2^. The monolithic CMOS device, fabricated in a standard 2-poly 4-metal 0.35-μm process, has an array of 32 × 32 electrochemical detectors (Fig. 6). The integration of the electrode array consists of platinum electrodes and SU-8 insulation as discussed in Section III.

**Fig. 5.**
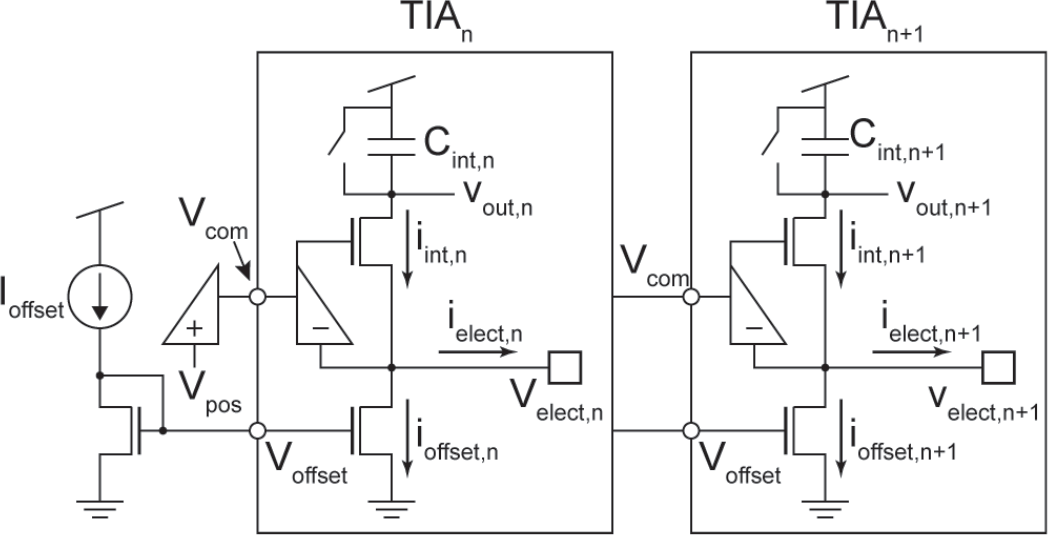
Half-shared OPA and TIA schematic. The non-inverting half of the OPA is shared among multiple TIAs. Each TIA has a dedicated electrode, integration capacitor, and current mirror generating I_offset_.

**Fig. 6.**
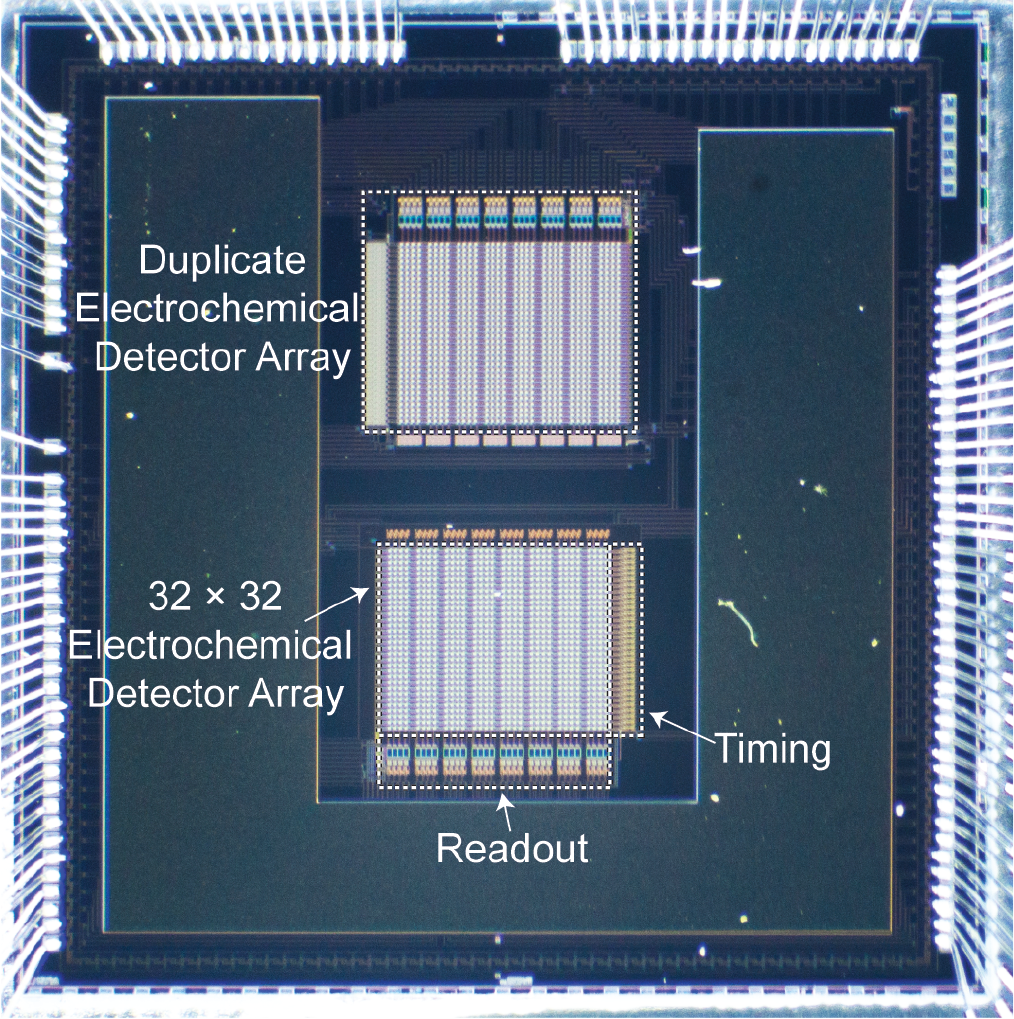
Photograph of the CMOS chip and the electrochemical detector arrays. To operate the array, two clocks are applied to the timing circuitry. The readout circuitry consists of the 32-to-1 multiplexers and output buffers to drive off-chip ADCs. A duplicate array is on the chip for characterization and testing.

As described in [11], the entire array is operated through the application of two clock signals to the timing circuitry which will generate various clock signals required to achieve correlated double sampling, resetting of the integration capacitor in each TIA, and time-division multiplexing. To condense the output of the 1024 electrochemical detectors to 32 parallel outputs on a column basis, time-division multiplexing is used. The time-division multiplexing staggers the readout period, as well as the integration period, of the 32 rows in a column to minimize integration deadtime of the array.

## III. CYCLIC VOLTAMMETRY ON AN ELECTRODE ARRAY

The goal of this study is to perform parallel CV on an electrode array to characterize the quality of individual electrodes. This section will discuss the integration of the electrode array onto the CMOS chip, how cyclic voltammograms reveal electrode characteristics, and the parallel CV measurements from an array of electrodes.

### A. Monolithic Integration of the Electrode Array

It is common for semiconductor foundries to fabricate CMOS chips with an aluminum-copper alloy as the top metal layer. However, aluminum is highly reactive to electrolytic solutions and introduces significant offsets and shot noise to electrochemical recordings. To prepare the device for electrochemical measurements, post-CMOS processing is used to integrate polarizable electrode materials with low reactivity, such as platinum or gold, onto the CMOS chip.

To integrate the platinum electrode array onto the CMOS chip, photolithography is used to perform a lift-off process. The chips are processed individually after they have been attached to a coverslip (25 mm × 25 mm) to ease the handling of the small devices (5 mm × 5 mm). After the chips have been attached to the coverslips, they are cleaned using acetone, isopropanol, and deionized water. For the lift-off process, a sacrificial layer of photoresist is created before metal deposition. This layer is created by spin coating, exposing, and developing a negative photoresist, NR9-1500PY, on the surface of the chip, removing photoresist where the electrodes will be patterned. The electrode materials are then deposited onto the processed chips, starting with the deposition of 20 nm of Ti and then 200 nm of Pt through sputtering (AJA Six-Gun Sputtering System, AJA International Inc., N Scituate, MA). After the metal is deposited onto the chip’s surface, the sacrificial layer of photoresist is removed by rinsing the sample with acetone. Removal of the sacrificial layer leaves only the desired electrodes on top of the amplifier array.

To prepare the monolithic CMOS device for electrochemical applications such as single-cell amperometry, a biocompatible epoxy-based photoresist, SU-8 3010, is used to provide isolation between the electrodes in the electrochemical detector array. After the lift-off process has been completed, the chips are again cleaned with acetone, isopropanol, and deionized water. The SU-8 photoresist is then spin coated, exposed, and developed, leaving isolating wells atop the electrodes with a size of 15 μm × 15 μm. The photoresist that remains on the surface of the monolithic CMOS device is approximately 20 – 30 μm thick.

### B. Parallel Cyclic Voltammetry

CV is an electrochemical measurement technique wherein the current is measured and plotted against the sweeping potential difference of the working electrode and the reference electrode, resulting in a cyclic voltammogram (Fig. 7). Throughout the electrochemical detector array, the potential of the integrated electrodes sweep from 1.2 – 2.0 V over a symmetric 10 s period while simultaneously measuring the current resulting from reactions occurring at the electrode’s surface. Phosphate-buffered saline (PBS) (2mL) is placed on the detector array and an Ag|AgCl reference electrode, connected to 1.2 V, is placed in the electrolytic solution to form the CV setup. As the potential of the integrated electrodes sweep from 1.2 – 2.0 V against the virtual ground of the Ag|AgCl electrode that is held at 1.2 V, the resulting applied potential in the electrolytic solution sweeps from 0 – 0.8 V at a rate of 10 s (Fig. 7a), resulting in a CV scan rate of 80 mV/s. To increase the dynamic range for the CV measurements, the array is run at a sampling rate of 20 kHz. The time domain current measurements from a representative electrochemical detector (Fig. 7b), comprised of a bipolar TIA and its platinum electrode, are done with and without PBS on the array. When there is no PBS on the array, the voltammogram in grey (Fig. 7c) shows a response that is predominantly resistive. After placing PBS on the array, the effects of the double-layer capacitance can be seen. This double-layer capacitance introduces a component to the measured current that reacts to change in the applied potential’s slope (Fig. 7b). When dealing with a purely capacitive system, the capacitance can be found by measuring the current that results from the applied potential’s slope through the relationship C = I/(dV/dt). However, the presented system has both resistive and capacitive impedance. To remove the influence of current due to resistive impedance, the change in current due to both the positive and negative slopes of the applied potential is examined. To maintain an objectiveness, 400 mV is chosen as a standard to review the change in current across the electrode array. Knowing the change in current at 400 mV and the symmetric slope of the applied potential, we can estimate the capacitance using C = ΔI/(2×dV/dt). From Fig. 7, the change in current at 400 mV is 104.8 pA and the slope of the applied potential is 80 mV/s. For this electrochemical detector the electrode’s double-layer capacitance is 655 pF.

**Fig. 7.**
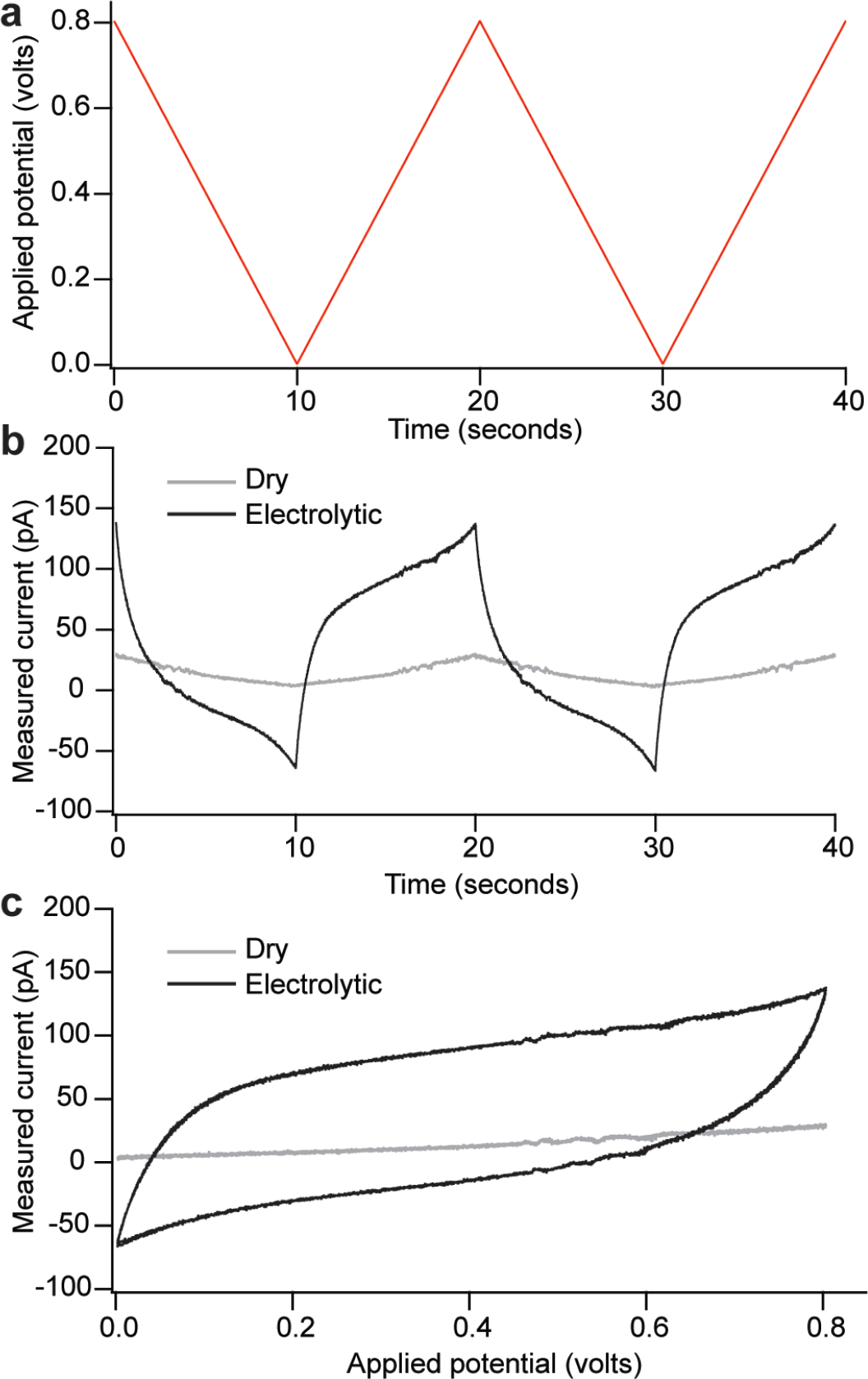
CV measurement from one TIA electrode. (a) Applied potential vs the reference electrode in the electrolytic solution. (b) Time domain current measurement from a TIA when the device is dry and when PBS is put on the array. (c) Voltammogram of the measured current versus the applied potential. The dry measurement is nearly resistive and the electrolytic measurement reveals the double-layer capacitance of the electrode.

To study the quality of the monolithically integrated electrodes, parallel CV is performed to characterize individual electrodes. For the designed electrochemical detector array the input V_pos_ is shared across the array, enabling simultaneous CV measurements. When measuring from the array of electrodes the setup is as previously described, the integrated electrode is swept from 1.2 – 2.0 V, the reference electrode is held at 1.2 V in 2 mL of PBS, and the sampling rate of each amplifier is 20 kHz. The resulting voltammograms from one parallel CV recording are shown in Fig. 8, illustrating various electrode qualities prior to any treatment. Reviewing electrodes 1 – 4, various levels of response are shown. In this group, electrode 4 is an outlier, appearing almost resistive in its response, thus indicating a weak interface between the electrode and the electrolyte.

**Fig. 8.**
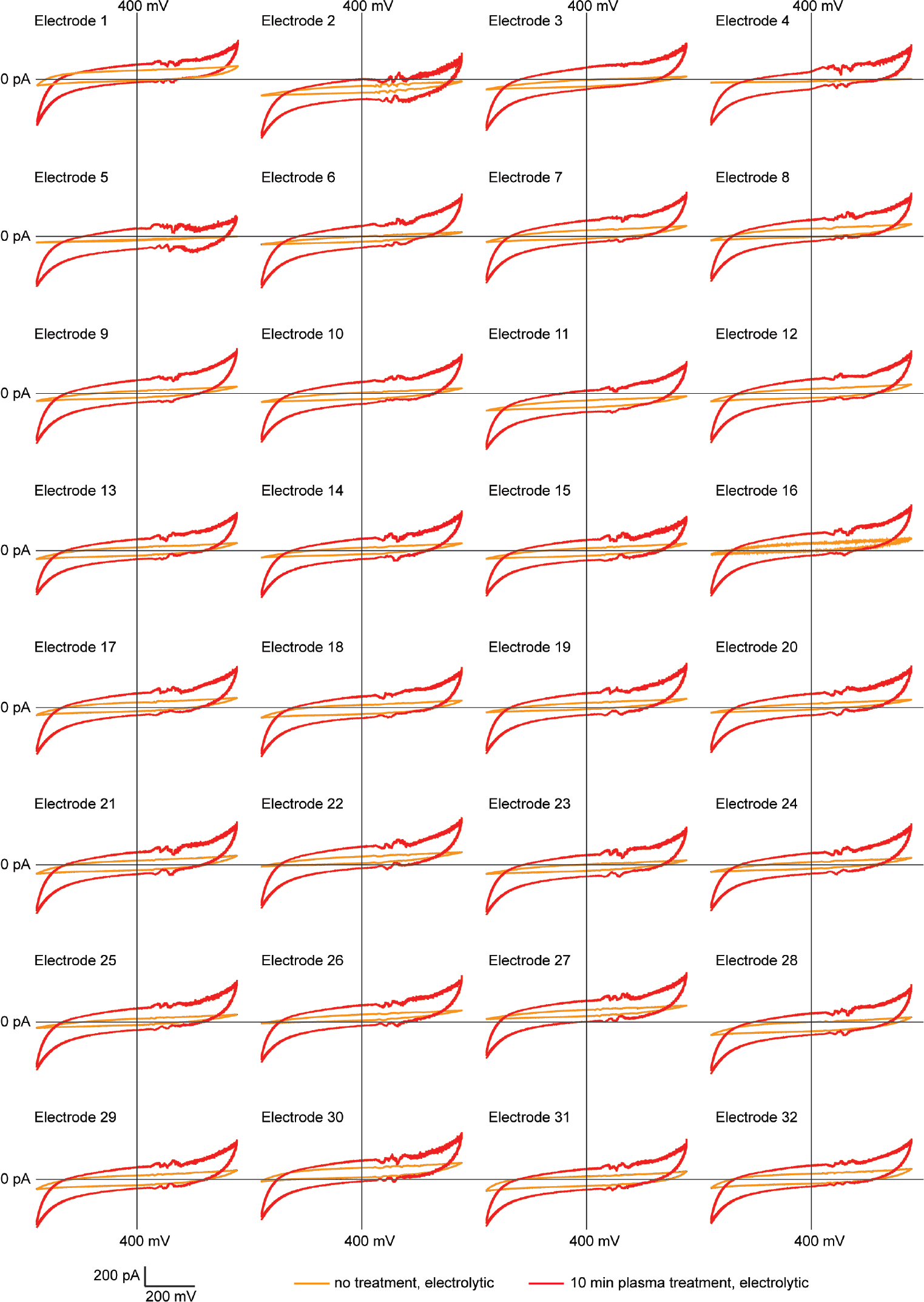
CV recordings from an electrode array. The electrodes are initially contaminated and exhibit reduced sensitivity as shown in the orange lines. After plasma treatment, the sensitivity of the electrode array is increased consistently throughout the array.

## IV. PARALLEL ELECTRODE ARRAY CHARACTERIZATION

### A. Electrode Surface Cleaning

After post-CMOS processing, cleaning of the electrode surface is required to ensure that the surface is pristine for use in electrochemical detections because contamination of the electrode’s surface reduces its effectiveness for electrochemical measurements. An electrode’s surface can be contaminated during the photolithography processes as well as through repeated use of the electrode for electrochemical measurements. In post-CMOS processing, the last step to create an insulation layer using SU-8 has been reported to often leave a thin film on the electrodes after development [7], [20]. Also, molecules, such as dopamine, can also adsorb onto the surface of the electrode [21], [22] after an electrochemical measurement and reduce the sensitivity of the electrode [23]. To clean the impurities throughout the electrode array, a plasma cleaner (PDC-32G, Harrick Plasma, Ithaca, NY) is used to remove organic material from the surface of the electrodes using ionized gas.

### B. Electrode Array Characterization Using Cyclic Voltammograms

The levels of contamination of an electrode array can be monitored by measuring the double-layer capacitance across the array through parallel CV recordings. As the area of the electrode’s surface is contaminated, the effective area that creates an interface between the electrode and the electrolyte, the double-layer capacitance, is reduced. Modeling the double-layer capacitance as C = εA/d, the reduction of the electrode-electrolytic interface area (A) is directly related to the reduction of capacitance. The electrochemical detector array, which was previously used for several experiments involving the oxidization of dopamine, is used for electrode array characterization using CV (Fig. 8, shown in orange). This set of voltammograms is used to calculate the double layer capacitance formed at each electrode-electrolytic interface. The average capacitance of the electrode array is 466 pF with a standard deviation of 143 pF. According to this analysis, electrodes 4 and 5 are found to have the most contamination with measured capacitances of 7.46 and 54.7 pF respectively. Conversely, electrodes 1 and 32 exhibit the lowest contaminations across the array with measured capacitances of 655 pF and 642 pF respectively. To remove the contaminants, the electrode array is given a 10-minute plasma treatment, and the response of the electrodes is re-characterized (Fig. 8, shown in red). The mean capacitance of the array after plasma treatment is 1.36 nF with a standard deviation of 84.5 pF. Electrodes 4 and 5 exhibit a capacitance of 1.17 and 1.34 nF respectively after treatment. The increased mean capacitance reveals that the electrodes could have been previously contaminated and plasma treatment improved the surface condition, increasing the area of the electrode-electrolyte interface and thus the double-layer capacitance.

## V. CONCLUSION

In this paper, we study the quality of an electrode array through parallel CV measurements, as well as present the design and performance of a bipolar capacitive TIA used in a 1024-channel monolithic CMOS device for high-throughput electrochemical recordings. The presented capacitive TIA is capable of measuring bipolar current from oxidation and reduction reactions through the introduction of a DC offset, I_offset_. The bipolar capabilities of the capacitive TIA are demonstrated by measuring alternating current that is injected into the electrode node. Integration of the bipolar capacitive TIA for large-scale parallel electrochemical recordings in a 1024-channel CMOS device is achieved through the use of the half-shared OPA structure. Through post-CMOS processing, the platinum electrode array and the SU-8 insulation is monolithically integrated on the CMOS device. To examine the quality of the electrode array after repeated use for electrochemical recordings, parallel CV is performed before and after plasma treatment. Before the array receives plasma treatment, some electrodes exhibit significant contamination, resulting in nearly resistive CV responses with capacitances of 7.5 and 54.7 pF compared to the array mean of 466 pF. The contaminations on the electrode surface, such as adsorbed molecules or a thin film of photoresist, reduce the effective area for the electrode-electrolyte interface and thus the double-layer capacitance. After plasma treatment of the electrode array, the mean capacitance of the array became 1.36 nF, indicating that the surface of the electrodes could have been contaminated and plasma treatment improved the surface of the electrode, thus increasing the electrode-electrolyte interface and the double-layer capacitance. Using parallel CV, characterization of the 1024 electrode array can be accelerated and conducted within minutes before it is used for analytical purposes.

## ACKNOWLEDGEMENTS

The authors thank Edward Dein for his technical assistance in the clean room. Post-CMOS processing was carried out in Advanced Microfabrication Facility (AMPAC) at the University of Central Florida.

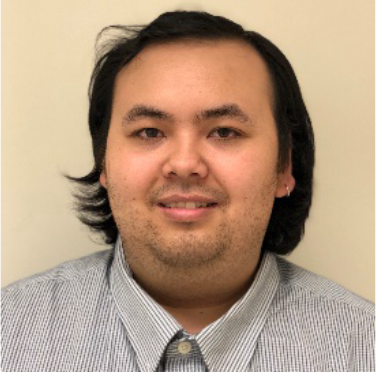

**Kevin A. White** received a B.S. degree in electrical engineering from the University of Central Florida, Orlando, FL, in 2016. He is a research assistant in Dr. Brian N. Kim’s research lab at the Burnett School of Biomedical Science since 2016. His research interests include circuit design, digital signal processing, post-CMOS processing, and brain-machine interfacing. He received a travel award to attend the Biophysical Society’s 61^st^ Annual Meeting in 2017.

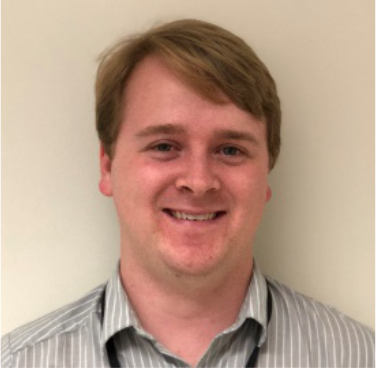

**Geoffrey Mulberry** was born in Cocoa Beach, Florida in March of 1995. He received a bachelor’s of science in electrical engineering from the University of Central Florida (UCF) in Orlando, Florida in 2017. He is currently a graduate student seeking a Ph.D. in electrical engineering from UCF. His main research interests are the development of medical diagnostic systems relying on electrical and mechanical methods, and miniaturization of such systems.

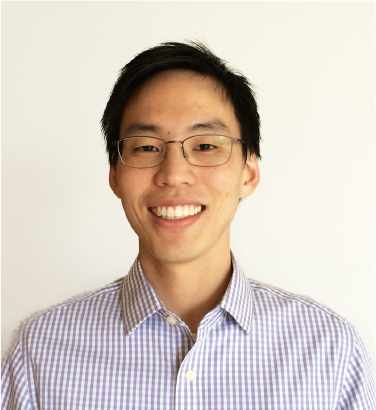

**Brian N. Kim** received the B.S. degree (with distinction) in Electronics and Computer Engineering from Hanyang University, Seoul, South Korea, and the Ph.D. degree in Biophysics from Cornell University, Ithaca, NY, in 2008 and 2013, respectively. Currently, he is Assistant Professor of Electrical and Computer Engineering at University of Central Florida, Orlando, FL. From 2014 to 2015, he was Senior Electrical Engineer at Stratos Genomics Inc., Seattle, WA. From 2013 to 2014, he was Postdoctoral Fellow in Bioengineering at University of California, Berkeley, CA. His research interests include single-cell electrophysiology, brain-machine interface, and medical diagnostics.

## Notes

This work was supported by National Science Foundation under grant #1745364.

## REFERENCES

[1] N. Kasai, A. Shimada, T. Nyberg and K. Torimitsu, “An electrochemical sensor array and its application to real-time brain slice imaging,” Electron. Commun. Japan, vol. 92, no. 9, pp. 1–6, Sep. 2009.

[2] J. Rothe, O. Frey, A. Stettler, Y. Chen and A. Hierlemann, “Fully Integrated CMOS Microsystem for Electrochemical Measurements on 32 × 32 Working Electrodes at 90 Frames Per Second,” Anal. Chem., vol. 86, no. 13, pp. 6425–6432, Jul. 2014.

[3] D. L. Bellin, H. Sakhtah, Y. Zhang, A. Price-Whelan, L. E. P. Dietrich and K. L. Shepard, “Electrochemical camera chip for simultaneous imaging of multiple metabolites in biofilms,” Nat. Commun., vol. 7, p. 10535, Jan. 2016.

[4] A. Hassibi and T. H. Lee, “A Programmable 0.18-$\mu\hbox{m}$ CMOS Electrochemical Sensor Microarray for Biomolecular Detection,” IEEE Sens. J., vol. 6, no. 6, pp. 1380–1388, Dec. 2006.

[5] J. Guo, W. Ng, J. Yuan, S. Li and M. Chan, “A 200-Channel Area-Power-Efficient Chemical and Electrical Dual-Mode Acquisition IC for the Study of Neurodegenerative Diseases,” IEEE Trans. Biomed. Circuits Syst., 2016.

[6] B. N. Kim, A. D. Herbst, S. J. Kim, B. A. Minch and M. Lindau, “Parallel recording of neurotransmitters release from chromaffin cells using a 10×10 CMOS IC potentiostat array with on-chip working electrodes,” Biosens. Bioelectron., vol. 41, pp. 736–744, 2013.

[7] M. Huang, J. B. Delacruz, J. C. Ruelas, S. S. Rathore and M. Lindau, “Surface-modified CMOS IC electrochemical sensor array targeting single chromaffin cells for highly parallel amperometry measurements,” Pflügers Arch. - Eur. J. Physiol., vol. 470, no. 1, pp. 113–123, Jan. 2018.

[8] K. Niitsu, S. Ota, K. Gamo, H. Kondo, M. Hori and K. Nakazato, “Development of Microelectrode Arrays Using Electroless Plating for CMOS-Based Direct Counting of Bacterial and HeLa Cells,” IEEE Trans. Biomed. Circuits Syst., vol. 9, no. 5, pp. 607–619, Oct. 2015.

[9] C. I. Dorta-Quinones, M. Huang, J. C. Ruelas, J. Delacruz, A. B. Apsel, B. A. Minch and M. Lindau, “A Bidirectional-Current CMOS Potentiostat for Fast-Scan Cyclic Voltammetry Detector Arrays,” IEEE Trans. Biomed. Circuits Syst., pp. 1–10, 2018.

[10] H. Li, S. Parsnejad, E. Ashoori, C. Thompson, E. K. Purcell and A. J. Mason, “Ultracompact Microwatt CMOS Current Readout with Picoampere Noise and Kilohertz Bandwidth for Biosensor Arrays,” IEEE Trans. Biomed. Circuits Syst., vol. 12, no. 1, pp. 35–46, 2018.

[11] K. A. White, G. Mulberry, J. Smith, M. Lindau, B. Minch, K. Sugaya and B. N. Kim, “Single-Cell Recording of Vesicle Release from Human Neuroblastoma Cells using 1024-ch Monolithic CMOS Bioelectronics,” IEEE Trans. Biomed. Circuits Syst., 2018.

[12] G. Mulberry, K. A. White and B. N. Kim, “Analysis of Simple Half-Shared Transimpedance Amplifier for Picoampere Biosensor Measurements,” IEEE Trans. Biomed. Circuits Syst., pp. 1–1, 2019.

[13] K. A. White, G. Mulberry and B. N. Kim, “Rapid 1024-Pixel Electrochemical Imaging at 10,000 Frames Per Second Using Monolithic CMOS Sensor and Multifunctional Data Acquisition System,” IEEE Sens. J., vol. 18, no. 13, pp. 5507–5514, Jul. 2018.

[14] H. Xicoy, B. Wieringa and G. J. M. Martens, “The SH-SY5Y cell line in Parkinson’s disease research: a systematic review,” Mol. Neurodegener., vol. 12, no. 1, p. 10, Dec. 2017.

[15] S. J. Chinta and J. K. Andersen, “Dopaminergic neurons,” Int. J. Biochem. Cell Biol., vol. 37, no. 5, pp. 942–946, May 2005.

[16] Y. Dong, M. L. Heien, M. M. Maxson and A. G. Ewing, “Amperometric measurements of catecholamine release from single vesicles in MN9D cells,” J. Neurochem., vol. 107, no. 6, pp. 1589–1595, Dec. 2008.

[17] M. Criado, A. Gil, S. Viniegra and L. M. Gutiérrez, “A single amino acid near the C terminus of the synaptosomeassociated protein of 25 kDa (SNAP-25) is essential for exocytosis in chromaffin cells.,” Proc. Natl. Acad. Sci. U. S. A., vol. 96, no. 13, pp. 7256–61, Jun. 1999.

[18] Q. Fang, Y. Zhao, A. D. Herbst, B. N. Kim and M. Lindau, “Positively charged amino acids at the SNAP-25 C terminus determine fusion rates, fusion pore properties, and energetics of tight SNARE complex zippering.,” J. Neurosci., vol. 35, no. 7, pp. 3230–9, 2015.

[19] S. Ayers, K. D. Gillis, M. Lindau and B. A. Minch, “Design of a CMOS Potentiostat Circuit for Electrochemical Detector Arrays,” IEEE Trans. Circuits Syst. I Regul. Pap., vol. 54, no. 4, pp. 736–744, Apr. 2007.

[20] J. Yao, X. A. Liu and K. D. Gillis, “Two approaches for addressing electrochemical electrode arrays with reduced external connections,” Anal. Methods, vol. 7, no. 14, p. 5760, 2015.

[21] R. F. Lane and A. T. Hubbard, “Differential Double Pulse Voltammetry at Chemically Modified Platinum Electrodes for in vivo Determination of Catecholamines.”

[22] J. O. Zerbino and M. G. Sustersic, “Ellipsometric and Electrochemical Study of Dopamine Adsorbed on Gold Electrodes,” 2000.

[23] K. Berberian, K. Kisler, Q. Fang and M. Lindau, “Improved Surface-Patterned Platinum Microelectrodes for the Study of Exocytotic Events,” Anal. Chem., vol. 81, no. 21, pp. 8734–8740, Nov. 2009.

